# CrowdGO: machine learning and semantic similarity guided consensus Gene Ontology annotation

**DOI:** 10.1101/731596

**Authors:** Maarten J.M.F. Reijnders, Robert M. Waterhouse

## Abstract

**Background:** Characterising gene function for the ever-increasing number and diversity of species with annotated genomes relies almost entirely on computational prediction methods. These software are also numerous and diverse, each with different strengths and weaknesses as revealed through community benchmarking efforts. Meta-predictors that assess consensus and conflict from individual algorithms should deliver enhanced functional annotations.

**Results:** To exploit the benefits of meta-approaches, we developed CrowdGO, an open-source consensus-based Gene Ontology (GO) term meta-predictor that employs machine learning models with GO term semantic similarities and information contents. By re-evaluating each gene-term annotation, a consensus dataset is produced with high-scoring confident annotations and low-scoring rejected annotations. Applying CrowdGO to results from a deep learning-based, a sequence similarity-based, and two protein domain-based methods, delivers consensus annotations with improved precision and recall. Furthermore, using standard evaluation measures CrowdGO performance matches that of the community’s best performing individual methods.

**Conclusion:** CrowdGO offers a model-informed approach to leverage strengths of individual predictors and produce comprehensive and accurate gene functional annotations.

**Availability and Implementation:** CrowdGO is implemented in Python3, and is freely available from https://gitlab.com/mreijnders/CrowdGO, with a Snakemake workflow and pre-trained models.

## Introduction

New technologies and decreasing costs are enabling rapid accumulation of large quantities of genomic data. Experimental elucidation of biological functions of genomic elements such as protein-coding genes lags behind because it requires considerable additional efforts. Exploiting the potential of genomic data therefore relies on bioinformatics approaches to predict functions. The most comprehensive and widely-used model of describing gene function is the Gene Ontology (GO), a cornerstone for biology research (Ashburner *et al*., 2000; The Gene Ontology Consortium, 2019). Numerous computational tools have therefore been developed aiming to transfer functional information in the form of GO term annotations from functionally characterised macromolecules to those currently lacking assigned functions (Zhao *et al*., 2020; Makrodimitris *et al*., 2020). Many software focus on using protein sequences to predict function through sequence homology and/or domains and motifs (Koskinen *et al*., 2015; Lavezzo *et al*., 2016; Scheibenreif *et al*., 2019; Mitchell *et al*., 2019). Others use protein sequence features in combination with similarity based predictions, or make use of complementary data sources where available, such as protein structures or protein-protein interactions (Yang *et al*., 2015; Kulmanov *et al*., 2018; Kulmanov and Hoehndorf, 2020). The function prediction community responded to the proliferation of tools with the Critical Assessment of Functional Annotation (CAFA) initiative to evaluate and improve computational annotations of protein function (Zhou *et al*., 2019). The common platform to benchmark performance reveals different strengths and weaknesses of participating predictors using standardised metrics. These measures are used to assess results from participating software for the Molecular Function Ontology (MFO), the Biological Process Ontology (BPO), and the Cellular Component Ontology (CCO).

Leveraging the strengths of individual algorithms through a consensus-conflict assessing meta-predictor should offer a means to achieve improved annotations of protein function. To test this potential, we developed CrowdGO, a consensus-based GO term meta-predictor that employs machine learning models with GO term semantic similarities and information contents (IC) to produce enhanced functional annotations. By re-evaluating each gene-term annotation, a consensus dataset is produced with high-scoring confident annotations and low-scoring rejected annotations. We assess the performance of CrowdGO results compared with four input annotation sets from a deep learning-based, a sequence similarity-based, and two protein domain-based methods. We also examine the effects of CrowdGO on reclassifying true and false positive and negative annotations. Using standard evaluation measures with the community benchmarking datasets, we compare CrowdGO results with those from CAFA3. Finally, we compare existing GO term annotations for several model and non-model species with those predicted using CrowdGO. The assessments and comparisons show that CrowdGO offers a model-informed approach to leverage strengths of input predictors and produce comprehensive and accurate gene functional annotations.

## Results

### Training, testing, and benchmarking annotations

To assess the performance of CrowdGO we first built training, testing, and benchmarking GO annotation sets following best practices for evaluating machine learning models (Table S1) and CAFA3 benchmark generation guidelines (see Materials and Methods). The set of annotations for building the CrowdGO models was derived from all proteins added to the GO Annotation (GOA) UniProt database from 2018 (version 162) to 2020 (version 198). The CAFA3 benchmarking dataset is older and consists of new GO term annotations added to the GOA UniProt database during 2017 as detailed in (Zhou et al., 2019). The V162-V198 training and testing annotation datasets have well-matched numbers of proteins per ontology, and a little over two thirds as many for MFO, about 10% more for BPO, and about double for CCO when compared with the CAFA3 benchmarking dataset (Table 1). The average numbers of leaf and parent-propagated GO terms per protein are generally similar across the three datasets, apart from slightly higher averages for CAFA3 MFO and nearly double the leaf annotations per protein for BPO in the V162-V198 datasets. Terms per protein and term ICs are also well-matched between the training and testing datasets, showing no significant differences in their distributions (Supplementary Figures S1 and S2). In the context of building a machine learning model, the V162-V198 dataset therefore offers a representative baseline to assess the performance of CrowdGO consensus results compared with annotations from individual input predictors. The CAFA3 benchmarking dataset is the standard in the field of protein function prediction, and thus offers a community reference to assess the performance of CrowdGO results compared with annotations from top-performing CAFA3 predictors.

**Table 1:** Average GO terms per protein for the V162-V198 training and testing datasets and the CAFA3 benchmarking dataset. Molecular Function (MFO), Biological Process (BPO), and Cellular Component (CCO) ontologies; leaves: GO terms with no child terms annotated for the same protein; propagated: after propagating parent GO term annotations.

| Dataset | Proteins | MFO |  | BPO |  | CCO |  |
| --- | --- | --- | --- | --- | --- | --- | --- |
|  |  | leaves | propagated | leaves | propagated | leaves | propagated |
| V162-V198 training | 335 MFO<br>988 BPO<br>1244 CCO | 1.3 | 7.4 | 6.8 | 32.0 | 4.7 | 13.7 |
| V162-V198 testing | 326 MFO<br>1000 BPO<br>1222 CCO | 1.3 | 7.3 | 6.1 | 29.4 | 4.6 | 13.5 |
| CAFA3 benchmarking | 454 MFO<br>848 BPO<br>639 CCO | 1.7 | 9.2 | 3.6 | 29.0 | 4.7 | 13.8 |

### CrowdGO consensus compared with input predictors

Next we used the V162-V198 training and testing annotation datasets to assess the performance of CrowdGO compared with annotations from four different individual input predictors comprising a deep learning-based, a sequence similarity-based, and two protein domain-based methods. Importantly, all datasets used to inform these predictors predate the annotated sequences in the V162-V198 dataset to ensure no overlaps, and the CrowdGO model was built with results from annotating the V162-V198 training dataset using these same predictors and datasets (see Materials and Methods). Precision-recall curves show the performance of individual results from DeepGOPlus (Kulmanov and Hoehndorf, 2020), FunFams (Scheibenreif et al., 2019), InterProScan (Jones et al., 2014), and Wei2GO (Reijnders, 2020) compared with results from applying CrowdGO to predictions from all four methods (Figure 1). For each of the ontologies, CrowdGO results show increased precision over annotations from the individual input predictors. Furthermore, CrowdGO presents a much smoother relationship between precision and recall across thresholds: the highest scoring annotations are the most reliable, and the lower scoring annotations steadily increase recall while reducing reliability. Evaluation metrics computed from the precision-recall curves (see Materials and Methods) show performance in terms of *F*_*max*_, the maximum protein-centric F-measure, *S*_*min*_ the Information Content (IC) semantic distance between true and predicted annotations, and area under the precision-recall curve (AUPR). CrowdGO results show improved *F*_*max*_, *S*_*min*_, and AUPR scores compared with annotations from each input predictor for each ontology (Table 2). The largest improvements across all evaluation metrics are with respect to the two protein domain-based methods, FunFams and InterProScan. The *F*_*max*_ improvements (Figure 1D) are largest for MFO, where the two next-best methods score similarly (DeepGOPlus and Wei2GO), but smaller for BPO and CCO where each has only one similarly-scoring next-best method. Assessing performance using the V162-V198 annotation dataset therefore demonstrates that the consensus approach implemented by CrowdGO successfully improves precision, recall, and *S*_*min*_ of gene-term annotations from each of the four individual input predictors.

**Figure 1:**
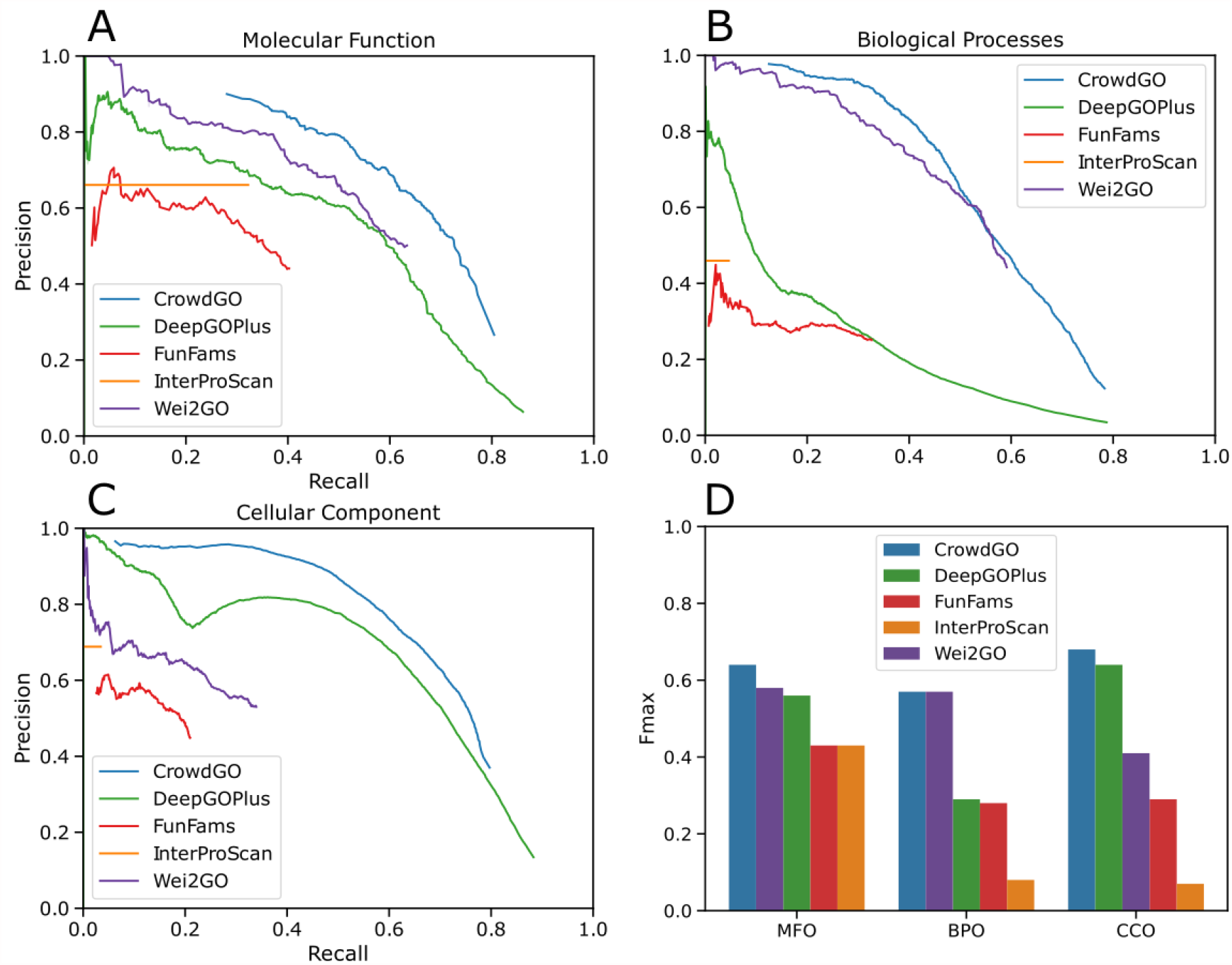
Precision-recall curves for GO term annotations from CrowdGO and four individual input predictors. Performance was assessed using the V162-V198 annotation dataset with threshold steps of probability scores of 0.01 for each predictor (except for InterProScan) for (A) the Molecular Function Ontology, (B) the Biological Process Ontology, and (C) the Cellular Component Ontology. *F*_*max*_ scores are shown in (D), sorted from highest to lowest scoring method for each ontology. Root terms are excluded from the evaluation. InterProScan predictions all have the same prediction probability (set to one), precision-recall curves are therefore presented as horizontal lines from zero recall to the actual recall.

**Table 2:**
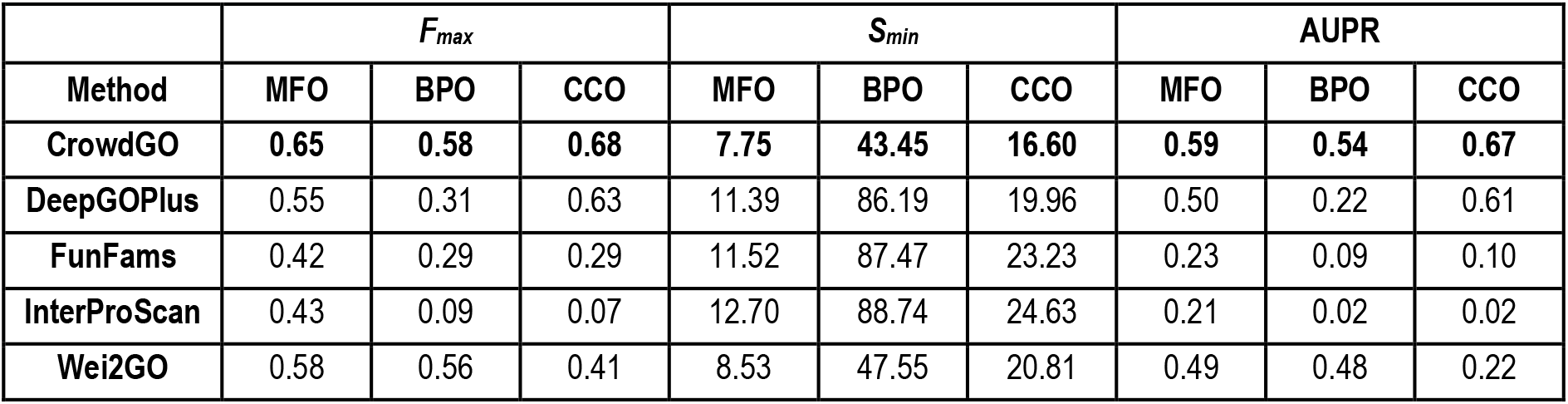
Evaluations of GO term annotations from CrowdGO and four individual input predictors. Molecular Function (MFO), Biological Process (BPO), and Cellular Component (CCO) ontologies; Highest *F*_*max*_ scores and AUPR values, and lowest *S*_*min*_ scores are in bold text.

| Method | $F_{max}$ | | | $S_{min}$ | | | AUPR | | |
| --- | --- | --- | --- | --- | --- | --- | --- | --- | --- |
|  | MFO | BPO | CCO | MFO | BPO | CCO | MFO | BPO | CCO |
| **CrowdGO** | **0.65** | **0.58** | **0.68** | **7.75** | **43.45** | **16.60** | **0.59** | **0.54** | **0.67** |
| DeepGOPlus | 0.55 | 0.31 | 0.63 | 11.39 | 86.19 | 19.96 | 0.50 | 0.22 | 0.61 |
| FunFams | 0.42 | 0.29 | 0.29 | 11.52 | 87.47 | 23.23 | 0.23 | 0.09 | 0.10 |
| InterProScan | 0.43 | 0.09 | 0.07 | 12.70 | 88.74 | 24.63 | 0.21 | 0.02 | 0.02 |
| Wei2GO | 0.58 | 0.56 | 0.41 | 8.53 | 47.55 | 20.81 | 0.49 | 0.48 | 0.22 |

### Impact of CrowdGO annotation re-evaluations

Because CrowdGO performs model-informed re-evaluations of scores from different input predictors, we next sought to quantify the impact of re-evaluations on the resulting annotations. Consensus predictors aim to leverage the information from multiple input methods to refine scoring and thereby improve overall calling of true positives and true negatives while decreasing false positives and false negatives. Thresholds based on the *F*_*max*_ achieved by each method using the V162-V198 annotation dataset were used to label true and false positives and negatives of the input annotations before applying CrowdGO and assessing the impacts of the re-evaluations (see Materials and Methods). By far the largest change for each of the three ontologies is the correction of false positives to true negatives (Figure 2). These changes dwarf the inverse, incorrect re-evaluations by CrowdGO that convert true negatives to false positives: correct re-evaluations are 11-56 times more numerous. The vast majority of true positives (75%-89%) and true negatives (84%-98%) are correctly affirmed, i.e. not re-classified, after applying CrowdGO. The consensus results are also able to recover false negatives from the input predictors and correct them to true positives, albeit for much smaller subsets. However, the consensus is somewhat conservative and therefore also incorrectly converts some true positives into false negatives. Comparing counts of true and false positives, CrowdGO consistently maximises true positive calls while minimising false positives compared to the results from the four different input predictors individually (Figure 2). The ratios are thus consistently improved over the individual inputs, and in several cases considerably so, e.g. for BPO from four false positives for each true positive for DeepGOPlus, and 7.5 false positives for each true positive for FunFams, to just one false positive for every three true positives. Assessing gene-term annotation re-evaluations using the V162-V198 annotation dataset therefore demonstrates that CrowdGO successfully decreases the calling of false positives while improving the overall calling of true positives and true negatives.

**Figure 2:**
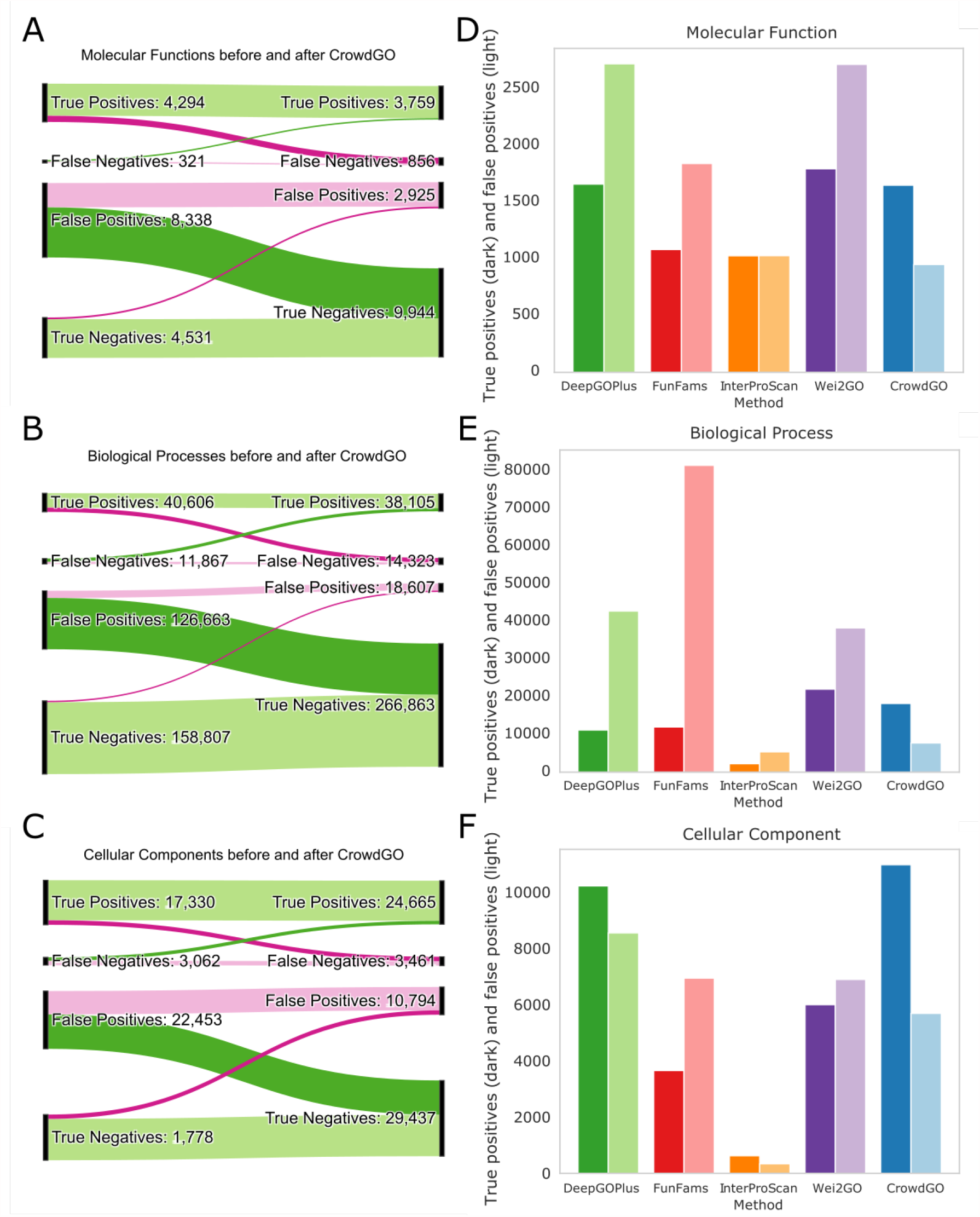
Impact of CrowdGO annotation re-evaluations on calling true and false positives and negatives. Sankey plots show the numbers of true or false positive or negative gene-term annotations before (left) and after (right) re-evaluation by CrowdGO for the three gene ontologies, (A) Molecular Function, (B) Biological Process, and (C) Cellular Component, for the V162-V198 testing dataset. Correct re-evaluations are shown in dark green (false positives to true negatives and false negatives to true positives), and incorrect re-evaluations are shown in dark pink (true positives to false negatives and true negatives to false positives). For unchanged annotations, correct affirmations are shown in light green (true positives and true negatives), and incorrect affirmations are shown in light pink (false positives and false negatives). Bar plots show for each method and CrowdGO (coloured as in Figure 1) counts of true positives (dark colours) and false positives (light colours) for the three gene ontologies, (D) Molecular Function, (E) Biological Process, and (F) Cellular Component, for the V162-V198 testing dataset.

### CrowdGO comparisons with CAFA3 methods

We next used the CAFA3 benchmarking dataset as a reference to compare CrowdGO results with annotations from baseline methods and predictors that took part in CAFA3 (Zhou *et al*., 2019). The four provided CrowdGO models (i.e. pre-trained on the complete V162-V198 dataset) were applied to GO term annotations produced by the four input predictors on the CAFA3 benchmarking dataset (see Materials and Methods). Performance summarised using the *F*_*max*_ scores (Table 3) shows that CrowdGO achieves *F*_*max*_ scores equal to the top-performing CAFA3 method for BPO, slightly lower than the best-scoring method but higher than the next-best performing methods for MFO, and slightly higher than the best CAFA3 methods for CCO. By this evaluation, the consensus approach applied to results from just four input predictors is able to produce annotations with precision and recall matching the *F*_*max*_ scores of the best-performing CAFA3 methods.

**Table 3:** *F*_*max*_ scores for CrowdGO and the baseline and top three performers from CAFA3. Molecular Function (MFO), Biological Process (BPO), and Cellular Component (CCO) ontologies; Highest *F*_*max*_ scores are in bold text.

| Method | $F_{max}$ | | |
| --- | --- | --- | --- |
|  | MFO | BPO | CCO |
| CAFA3 #1 | **0.62** | **0.40** | 0.61 |
| CAFA3 #2 | 0.54 | 0.39 | 0.61 |
| CAFA3 #3 | 0.53 | 0.38 | 0.60 |
| CAFA3 Naïve | 0.33 | 0.26 | 0.54 |
| CAFA3 BLAST | 0.42 | 0.26 | 0.46 |
| CrowdGOFull | 0.57 | **0.40** | **0.62** |
| CrowdGOLight | 0.57 | **0.40** | **0.62** |
| CrowdGOFull-SwissProt | 0.57 | 0.38 | 0.61 |
| CrowdGOLight-SwissProt | 0.57 | 0.39 | 0.60 |

### CrowdGO applied to model and non-model species

Having established that CrowdGO delivers consensus annotations with improved precision and recall matching the community’s best performing methods, we next assessed the ability of CrowdGO to functionally annotate complete proteomes of taxonomically diverse species. CrowdGO was applied to the full sets of protein-coding genes from a selection of animals, plants, fungi, and bacteria (see Materials and Methods). The confident consensus results were compared to existing GO term annotations from UniProt with respect to numbers of annotated proteins, and distributions of numbers of GO Slim terms per protein and total leaf+parents GO term informativeness (Figure 3, Supplementary Figure S3). Firstly, across the ontologies total numbers of annotated proteins increase for all 12 species, with larger increases for non-model species that have generally less comprehensive experimental data to support functional annotations. Secondly, the breadth of functional annotations also increases, with medians increasing by one to four GO Slim terms per protein compared to the TrEMBL or UniProt annotation datasets. Finally, the depth of functional annotations in terms of total informativeness of annotations per protein also increases, most notably for non-model species and compared to TrEMBL annotation datasets of model species. For model species with subsets of electronically inferred TrEMBL and manually reviewed SwissProt annotations such as *Arabidopsis thaliana* (Figure 3A) and *Drosophila melanogaster* (Figure 3B), the SwissProt annotations show more GO Slim terms and higher total informativeness per protein, albeit with fewer annotated proteins. In both these species, annotation breadth and depth of the high-quality SwissProt subsets more closely resemble those of CrowdGO than TrEMBL annotations, despite CrowdGO covering the largest numbers of proteins. Thus CrowdGO dramatically increases proteome coverage compared to SwissProt, while adding more terms, and more informative terms, compared to TrEMBL. The model bacterium, *Escherichia coli* (Figure 3C), and yeast, *Saccharomyces cerevisiae* (Figure 3D), are both fully SwissProt-reviewed, so CrowdGO only marginally increases proteome coverage. Compared with these manually curated gold standards, the automated predictions inevitably result in decreased annotation breadth and depth. The non-model species on the other hand have few or no SwissProt-reviewed annotations, thus their full UniProt annotations (TrEMBL+SwissProt) are used for comparisons, where CrowdGO results show improved proteome coverage with more GO Slim terms and higher total informativeness per protein. Notably, breadth and depth of CrowdGO annotations of the proteomes of these non-model species more closely resemble those of the high-quality SwissProt annotations for the model species than their own UniProt annotations.

**Figure 3:**
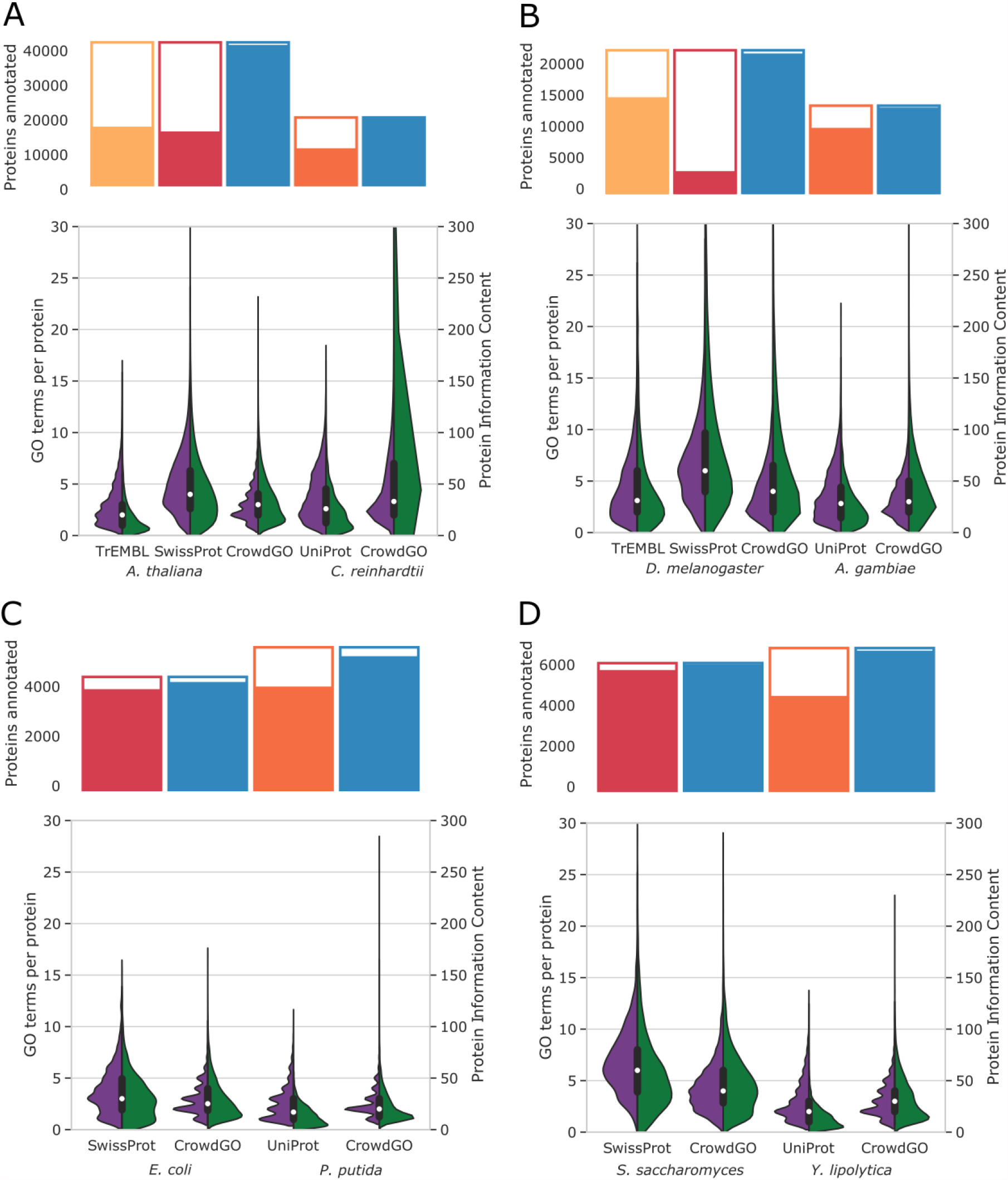
Comparisons of whole-proteome CrowdGO annotations and existing UniProt annotations. CrowdGO consensus annotation results were compared with existing Gene Ontology (GO) term annotations from UniProt, and with the subsets of manually curated SwissProt (where available) and automatically inferred TrEMBL annotations, for representative taxa of (A) plants and algae, (B) insects, (C) bacteria, and (D) fungi. For each species traditionally considered as a model species (*A. thaliana, D. melanogaster, E. coli*, and *S. cerevisiae*) a non-model species is used for comparison (*C. reinhardtii, A. gambiae, P. putida*, and *Y. lipolytica*). Bars at the top of each panel show the total numbers of proteins annotated with at least one GO term for each respective annotation dataset, with white-filled areas showing the remaining proteins with no annotations. The most general annotations are excluded from the total numbers of proteins annotated by counting only GO terms with at least two parents. Split violin plots show the distributions of the numbers of GO Slim terms annotated per protein (purple, left), and summed leaf+parents information content (IC) per protein (green, right). Y-axes are limited to a maximum of value of 300 for total protein IC distributions. The boxplots show the medians and 1.5 times the interquartile range for the numbers of GO Slim terms annotated per protein.

## Discussion

Here we explore whether a consensus-based GO term meta-predictor that leverages the strengths of individual algorithms can achieve improved computational annotations of protein function. Our results demonstrate that enhanced sets of functional annotations can be produced by employing machine learning models with GO term semantic similarities and ICs to re-evaluate sets of input annotations. Assessing CrowdGO consensus annotations using a dataset derived from all new SwissProt proteins over a 2.5-year period shows improved precision, recall, and *S*_*min*_ over each of four individual input predictors. Examining the effects of annotation re-evaluations further demonstrates that CrowdGO successfully decreases the calling of false positives while improving the overall calling of true positives and true negatives. Using the CAFA3 benchmarking dataset shows that applying CrowdGO to results from just four input predictors is able to produce annotations with precision and recall matching the *F*_*max*_ scores of the best-performing CAFA3 methods. Annotating complete proteomes of a diverse selection of species demonstrates that, while consensus predictions cannot match gold standard models like *E. coli* and *S. cerevisiae*, they can substantially improve coverage, breadth, and depth of functional annotations for a wide range of organisms. Using a model-informed meta-predictor approach, CrowdGO is able to re-evaluate results from individual predictors and produce comprehensive and accurate functional annotations.

We employed as input for model training just four predictors: the deep learning-based DeepGOPlus method, the sequence similarity-based Wei2GO approach, as well as the InterProScan and FunFams protein domain-based methods. DeepGOPlus and Wei2GO were selected because they are both entirely open-source and easily implementable. FunFams is the only method of these four that participated in CAFA3, where it performed well (amongst the top 10 for all three ontologies). It is also open-source, although to obtain annotation scores we had to implement scoring based on details provided in (Rentzsch and Orengo, 2013). InterProScan is freely available and widely used for annotating protein domains, with quality-assured InterPro2GO domain-to-term mapping generated through manual curation (Burge *et al*., 2012). We did not include the overall CAFA3 best performer, GOLabeler (You *et al*., 2018), because without an available open-source implementation it is not possible to train the machine learning classifiers that CrowdGO relies on to perform the consensus re-evaluations. Other CAFA3 top-performers offer only in-house or web-server implementations, and thus currently cannot be used with CrowdGO. Future open-source implementations of these and other newer methods would allow for their integration into CrowdGO through providing additional sets of pre-trained models.

Our evaluations indicate that CrowdGO performance in terms of *F*_*max*_ scores is primarily driven by one or two of the input predictors, namely DeepGOPlus for CCO, Wei2GO for BPO, and both these methods for MFO (Figure 1D). The two domain-based methods contribute less overall, particularly for CCO and BPO predictions, but with more substantial contributions for MFO, where CrowdGO improves the most over the next-best methods. A simple consensus approach such as retaining only annotations from the best-performing input method that are also supported by at least one other method could still reject true positives. Instead, the CrowdGO consensus approach performs model-informed re-evaluations of the scores of all annotations based on predictions from all inputs, enabling it to improve on the best input method, especially for MFO. Therefore, assuming that good models can be built with additional methods, the inclusion of more input annotation sets could lead to enhanced performance improvements. Furthermore, the re-evaluations also produce scores that more smoothly balance precision and recall allowing users to select larger sets of less confident annotations or to focus on only the highest scoring most reliable annotations.

The CrowdGO models were built using an Adaptive Boosting (Freund and Schapire, 1997) machine learning model, which aims to combine a set of weak classifiers into a weighted sum representing the boosted strong classifier. Future implementations of CrowdGO, especially with the inclusion of additional input predictors, might benefit from developing additional models using eXtreme Gradient Boosting, a popular alternative that is a powerful algorithm for supervised learning tasks (Chen and Guestrin, 2016). Given the variety of formulations used for measuring semantic similarities (Mazandu *et al*., 2016; Pesquita, 2017), and the recognition of the power of using ontologies to compute similarity and incorporate them in machine learning methods (Kulmanov *et al*., 2020), future developments to CrowdGO could also offer users models that are based on alternatives to the currently implemented Lin’s measure. Results from applying the model-informed approach of CrowdGO demonstrate that a consensus meta-predictor can improve protein functional annotations, as well as presenting opportunities for future models built with new training data, incorporating new predictors, and alternative boosting, to further enhance performance.

## Materials and Methods

### The CrowdGO algorithm

CrowdGO examines similarities between annotations from different GO term predictors, using ICs, semantic similarities, and a machine learning model to re-evaluate the input annotations and produce consensus results. Input for CrowdGO consists of (i) gene-term annotations from individual GO term predictors; (ii) the GO Annotation (GOA) database for UniProt (The UniProt Consortium, 2019) proteins; and (iii) a pre-trained Adaptive Boosting (AdaBoost) (Freund and Schapire, 1997) machine learning model. Annotations must be provided as a text file in tabular format consisting of four columns: predictor name, protein identifier, GO term identifier, and probability score. The GOA is provided with CrowdGO or can be downloaded from UniProt. Pre-trained AdaBoost models are provided, and new models can be trained using CrowdGO accessory functions.

The IC for each GO term from the input datasets is computed as the relative frequency of a GO term *i* compared to the total number of GO terms in the UniProt GOA database:

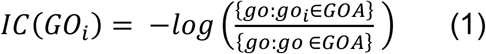

The ICs are then used to compute semantic similarities across all pairs of GO terms assigned to each protein by any of the input methods. GO term semantic similarities employ the GO’s directed acyclic graph (DAG) to define a metric representing the closeness in meaning between pairs of terms (Pesquita, 2017). It is computed here with *S* being the subset of GO terms shared between terms *i* and *j* after propagating up the GO DAG using the ‘is_a’ and ‘part_of’ relations, using the formula proposed by Lin (Lin, 1998):

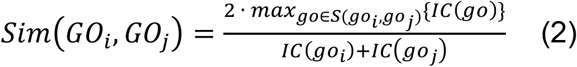

Gene-term annotations from all input methods are then compared using these similarity scores. For each input dataset, support for a given annotation is collected in the form of a list of the most similar GO term from each of the other methods, requiring a semantic similarity score of 0.5 or higher. Each gene-term annotation is therefore associated with the provided predictor method and probability score, and the GO term’s IC. When other methods predict the same or similar GO terms then the support list for each annotation comprises each of these additional GO terms, the provided predictor method and probability score, the computed IC, and the semantic similarity score with the GO term of the annotation they support. The support lists for each annotation are then used as features in pre-trained AdaBoost machine learning models. Based on these features, the model returns a probability score for each unique annotation from all input datasets. These are used to re-evaluate each annotation and produce the consensus dataset split into confident annotations with scores of 0.5 or higher, and the remaining rejected annotations.

CrowdGO provides four pre-trained models in pickle serialized file format: CrowdGOFull, CrowdGOLight, CrowdGOFull-SwissProt, and CrowdGOLight-SwissProt. CrowdGOFull is trained using results from four predictors: DeepGOPlus (Kulmanov and Hoehndorf, 2020); Wei2GO (Reijnders, 2020); InterProScan (Jones *et al*., 2014); and FunFams (Scheibenreif *et al*., 2019). CrowdGOLight excludes InterProScan, and the ‘-SwissProt’ models use input from Wei2GO based only on SwissProt proteins rather than the full UniProt set. The pre-trained AdaBoost models are built by supervised learning with labelled training data consisting of annotations from SwissProt (see below). The scikit-learn (Pedregosa *et al*., 2011) AdaBoost models are trained using a decision tree classifier with a depth of 2 as base estimator, a maximum of 100,000 iterations, a learning rate of 1, and the SAMME algorithm. AdaBoost scores are calibrated using the scikit-learn CalibratedClassifierCV package with 10-fold cross validation.

### Proteins and gene ontology annotations

The V162-V198 training, testing, and CAFA3 benchmarking GO annotation sets were built following best practices for defining datasets for evaluating machine learning models (Table S1) and CAFA3 benchmark generation guidelines.

#### The V162-V198 dataset

The set of annotations for building the AdaBoost models was derived from all proteins added to the GOA UniProt database from version 162 (17-01-2018) to version 198 (17-06-2020). Only annotations for proteins from the manually annotated SwissProt subset of UniProt, and with evidence codes EXP, IDA, IPI, IMP, IGI, IEP, TAS, or IC were retained, and proteins included in the CAFA3 benchmarking datasets were removed. If the only MFO term for a protein was ‘GO:0005515’ (protein binding), this was removed from both the training and test sets (326 out of 967 proteins), in line with CAFA3 benchmark generation guidelines. Finally, all parent GO terms from the GO DAG (release 2016-11-12) were propagated onto each of the annotations using the ‘is_a’ and ‘part_of’ relationships. This resulted in 661 proteins annotated with 4,477 MFO terms (397 unique, 3.3 % of MFO), 1,988 with 61,043 BPO terms (2,800 unique, 9% of BPO), and 2,466 with 33,564 CCO terms (407 unique, 9.3% of CCO). This complete V162-V198 dataset used to train the AdaBoost models is provided with CrowdGO. In order to compare CrowdGO results with annotations from individual input predictors, this dataset was split by randomly assigning 50% of proteins into a V162-V198 training subset and a V162-V198 testing subset.

#### The CAFA3 benchmarking dataset

CAFA benchmarking datasets of proteins and their GO terms rely on annotation growth over time and are generated by selecting all proteins that have gained experimental annotations during a set period. To assess the performance of CrowdGO results compared with annotations from the baseline and top-performing CAFA3 predictors, the CAFA3 no-knowledge (type1) benchmarking dataset was used, which consists exclusively of proteins without any prior annotations. This CAFA3 benchmarking dataset consists of new GO term annotations added to the GOA UniProt database from 13-02-2017 to 15-11-2017 as detailed in (Zhou *et al*., 2019).

#### Model and non-model species proteomes

CrowdGO was applied to complete proteomes (all protein-coding genes in the genome) for 12 model and non-model species to compare the results with their existing annotations. These, together with their SwissProt (where available) and TrEMBL annotations, were retrieved from the UniProt proteomes (downloaded 18-05-2020) for: *Anopheles gambiae* mosquito (UP000007062); *Arabidopsis thaliana* mouse-ear cress (UP000006548); *Candidatus Thorarchaeota archaeon* SMTZ1-45 (UP000070149); *Chlamydomonas reinhardtii* green alga (UP000006906); *Drosophila melanogaster* fruit fly (UP000000803) *Escherichia coli* K12 Gram-negative bacterium (UP000000625); *Homo sapiens* human (UP000005640); *Pan troglodytes* chimpanzee (UP000002277); *Pseudomonas putida* Gram-negative bacterium (UP000250299); *Saccharomyces cerevisiae* yeast (UP000002311); *Solanum lycopersicum* tomato (UP000004994); and *Yarrowia lipolytica* yeast (UP000256601).

### Running input predictors and CrowdGO

CrowdGO performance was assessed by comparing the consensus results with four input annotation sets from running: DeepGOPlus, Wei2GO, InterProScan, and FunFams, on the V162-V198 testing dataset. DeepGOPlus (GitHub 13-09-2020, using the DeepGOPlus 2016 dataset) was run with default parameters. DeepGOPlus and Wei2GO require DIAMOND (Buchfink *et al*., 2015) for protein sequence searches, and were run with version v0.9.11.112. Wei2GO and FunFams require HMMScan and HMMSearch functions from HMMER (Eddy, 2011), and were run using HMMER version 3.3.1. Wei2GO v1.0 was run with default parameters using sequences and annotations from UniProt release 2016-11 and profiles from Pfam-A (El-Gebali *et al*., 2019) version 30 (2016-05), i.e. with datasets that predate the sequences in the V162-V198 dataset to ensure no overlaps. InterProScan version 5.21.60 (2016-11) was run with default parameters and the --goterms option to obtain GO term annotations, using all default member databases as well as PANTHER protein families from version 10 (Mi *et al*., 2016), i.e. families that predate the sequences in the V162-V198 dataset. FunFams (cath-tools-genomescan GitHub 17-12-2019) was run with default parameters with CATH database version 4.1 (Dawson *et al*., 2017), which was based on Protein Data Bank (PDB) release 01-01-2015. InterProScan and FunFams do not provide probability scores for their predictions. For FunFams these were computed as detailed in (Rentzsch and Orengo, 2013), briefly: annotation frequency or normalised GO term occurrence count takes into account the likelihood of co-occurring with protein domains with other functions, so that each FunFams family is associated with a term-specific probability value. Probability scores for assignments from different InterProScan member databases are not directly comparable, so all InterProScan predictions were set to a score of 1. Finally, an AdaBoost model was built using results from annotating the V162-V198 training dataset using DeepGOPlus, Wei2GO, InterProScan, and FunFams. This AdaBoost model was then used to re-evaluate each gene-term annotation score and produce the new consensus dataset of annotations for the V162-V198 testing dataset. Performance was also compared with CAFA3 predictors, using the pre-trained AdaBoost models provided with CrowdGO to produce consensus annotations from results obtained by running DeepGOPlus, Wei2GO, InterProScan, and FunFams on the CAFA3 benchmarking dataset.

### Performance evaluation metrics

Performance was assessed using the V162-V198 training and testing datasets to compare with the four input methods, and using the CAFA3 no-knowledge benchmarking dataset to compare with CAFA3 predictors. For all evaluation metrics, threshold steps of probability scores 0f 0.01 were used for outputs of each predictor (except for InterProScan due to uniform probability score of 1), with Wei2GO probability scores first scaled from zero to one. The F1-score summarises performance as the harmonic mean between precision and recall, with *F*_*max*_ as the highest F1-score across all thresholds (Friedberg and Radivojac, 2017). Considering GO term ICs is also important, as annotating general terms is less useful than predicting more specific functions. The *S*_*min*_ score measures the minimum semantic distance between the predicted and real annotations across all thresholds (Clark and Radivojac, 2013). These standard benchmarking methods were used to define the evaluation metrics for precision, recall, *F*_*max*_, *S*_*min*_, and area under the precision-recall curve (AUPR) calculated as follows:

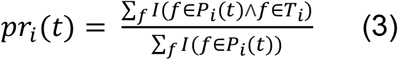

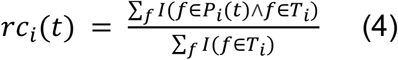

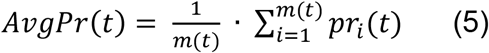

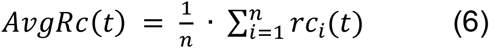

Where for each threshold (*t*) the precision (*pr*) and recall (*rc*) are calculated per protein (*i*), *f* is a GO term, *T*_*i*_ is the set of true terms annotated to protein _i_, and *P*_*i*_ the set of predicted terms annotated to protein _i_, *m*(*t*) is the number of proteins for which at least one term is predicted above the threshold, *n* is the total number of proteins, and *I* is an identity function which returns 1 if True and 0 if False. The maximum F1-score is then calculated, with thresholds ranging from 0 to 1 in steps of 0.01.

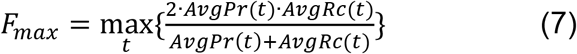

*S*_*min*_ is computed per protein based on the remaining uncertainty (*ru*), which is the IC of all GO terms *g* that comprise the subset of true terms annotated to the protein that were not predicted (False Negatives), and the missing information (*mi*), which is the IC of all terms *g* that comprise the subset of predicted terms that are not in the set of true terms annotated to the protein (False Positives).

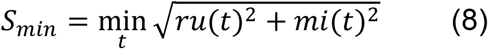

Where *ru* and *mi* are calculated as:

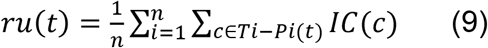

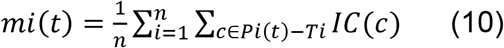

### Assessing impact of CrowdGO re-evaluations

To examine the effects of CrowdGO re-evaluations of the input annotation sets, correct and incorrect classifications of true and false positive and negative annotations were enumerated by comparing annotations before and after applying CrowdGO. Each gene-term annotation from the predictions of all input methods on the V162-V198 testing set was labelled true positive (tp), false positive (fp), true negative (tn), or false negative (fn). All thresholds used for assigning positive or negative labels were based on the *F*_*max*_ scores that each input method achieved on the V162-V198 testing dataset. Confident CrowdGO consensus annotations (scores of 0.5 or higher) were labelled positives and the rejected annotations were labelled negatives. The total numbers of correct (fn->tp; fp->tn) and incorrect (tp->fn; tn->fp) reclassifications, and of classification affirmations that were correct (tp->tp; tn->tn) or incorrect (fn->fn; fp->fp), were visualised using Sankey plots for each ontology.

### Annotating model and non-model species

CrowdGO was applied to annotate the proteomes of 12 species using default parameters and the provided CrowdGOFull AdaBoost model. FunFams was run using default parameters with version funfam-hmm3-v4_3_0.lib (based on PDB release 01-07-2019); InterProScan was run using default parameters and the --goterms option with version 5.45-80.0 (18-06-2020); Wei2GO was run using default parameters with UniProt release 06-2020 and Pfam version 33.0 (18-03-2020); and DeepGOPlus was run with default parameters using the GitHub repository downloaded on 01-06-2020. Consensus CrowdGO results were compared with existing annotations for the 12 species from SwissProt (where available) and TrEMBL databases from UniProt. All annotations from UniProt were included and compared to confident CrowdGO annotations (scores of 0.5 or higher). For both UniProt and CrowdGO datasets, all parent GO terms were propagated onto each gene-term annotation using the GO DAG (obo version 23-03-2020). To quantify the breadth of terms annotated to each protein, counts of terms per protein were calculated by summing all parent terms that are part of the GO slim ‘Generic GO Subset’ (downloaded 10-10-2020). GO term ICs computed as defined in equation (1) were used to quantify annotation depth by summing the IC of all terms (including parent terms) annotated to each protein. These measures of annotation breadth (counts of GO Slim terms per protein) and depth (summed IC of terms and their parents) were used to compare CrowdGO results with existing annotations for each species.

### CrowdGO implementation and availability

CrowdGO is available from https://gitlab.com/mreijnders/CrowdGO. The distribution includes DeepGOPlus (GitHub version 01-06-2020), FunFams (GitHub version 01-06-2020), and Wei2GO (GitLab version 15-11-2020). The user guide details instructions for also including InterProScan. Snakemake (Köster and Rahmann, 2012) workflows are provided to run each of the individual predictors, and to run CrowdGO to produce consensus annotations. CrowdGO can also be run directly with Python and the user-provided results from individual predictors, as detailed in the user guide. The distribution includes the four pre-trained AdaBoost models as well as supporting scripts to train new CrowdGO prediction models, e.g. when using other predictors to generate input annotations.

### CrowdGO software workflow

#### Step 1: read the user input arguments

**Function name:** readArguments()

**Function purpose:** reads the user-given CrowdGO input file, output directory, and model file (only when predicting GO terms)

CrowdGO input predictions are given in a tabular format using five columns for each prediction when training a model, and four columns for each prediction when predicting GO terms. Each row represents one prediction from one predictor: (1) name of one of the input predictors used in the trained model, (2) protein accession, (3) predicted GO term, (4) associated prediction score, (5) True or False label (only when training a model).

The output folder will contain the CrowdGO predictions and associated provenance when predicting GO terms, and will contain the trained model as a python pickled file and associated provenance when training a new model.

A model file is only given when predicting GO terms. Either one of the pre-trained models can be used, or a new model can be trained. It is highly recommended to use a model matching the ontology to be predicted, i.e. ‘bp.pkl’ for biological process terms, ‘mf.pkl’ for molecular function terms, and ‘cc.pkl’ for cellular component terms.

#### Step 2: read the input files

**Function name:** readInput()

**Function purpose:** reads the CrowdGO input files for training or predicting

This function reads the given input predictions from different predictors into a dictionary ‘predictionDictionary’.

#### Step 3: calculate GO relations according to the DAG

**Function name:** goSlim()

**Function purpose:** Creates several dictionaries containing direct and indirect GO parent and GO child relations for each GO term.

The data is parsed from the ‘goParents.tab’ and ‘goChildren.tab’ files in the CrowdGO data folder. A dictionary is made containing all direct and indirect parents of each GO term, a separate dictionary for all child terms of each GO term, and a dictionary containing all parent and child terms for each GO term. These dictionaries are later used to calculate similarities between GO terms predicted for a protein between the different input methods.

#### Step 4: parse the number of occurrences of each GO term in the GOA database

**Function name:** getGoCounts()

**Function purpose:** For each GO term its total occurrences in the GOA database are stored in a dictionary.

This data is parsed from the ‘goCounts.tab’ file in the CrowdGO data folder. A dictionary containing the counts for each GO term is returned, as well as a sum of the total GO count of the GOA database. These counts are later used to calculate the semantic similarity between GO terms for the same protein between different predictors.

#### Step 5: parse the namespace belonging to each GO term

**Function name:** getNameSpaces()

**Function purpose:** for each GO term its namespace, e.g. ‘biological_process’, is stored in a dictionary.

This data is parsed from the ‘nameSpaces.tab’ file in the CrowdGO data folder. GO namespaces are used to see if two GO terms are potentially related, and to calculate similarities between GO terms.

#### Step 6: calculate total GO terms belonging to a namespace

**Function name:** namespaceGOCount()

**Function purpose:** store the total occurrences of all GO terms belonging to a specific namespace in a dictionary.

This data is parsed from a dictionary of GO counts calculated in step 4, and the dictionary of namespaces parsed in step 5. These total counts for a namespace are later used in the calculation of similarities between GO terms.

#### Step 7: create a dictionary of similar GO terms predicted between methods for each protein

**Function name:** createClusters()

**Function purpose:** for every predicted GO term, find similar GO terms predicted by different methods and store this information in a dictionary.

For each row in the input file, i.e. for each prediction by each predictor, finds the parent GO terms from the other predictors that are most similar. Only GO terms with a semantic similarity of 0.5 or higher are considered. This results in each prediction containing the original GO term, as well as potentially the GO term from each other predictor that is the most similar. Additional information that is stored: the information content for each GO term, the prediction score for each GO term as given by the original prediction, the semantic similarity between each GO term from each different method, and the total amount of different input predictors that predicted a similar GO term. When training a model, and additional ‘True’ or ‘False’ label is stored as a ‘1’ or ‘0’ respectively, to indicate whether the GO term prediction is correct.

#### Step 8: train a model or predict GO terms using a model

**Function name:** model()

**Function purpose:** Either trains an AdaBoost model given the data created in step 7, or use an existing AdaBoost model to predict GO terms.

The features and labels (when applicable) are read from the files created in step 7. Output is written to the user specified output folder, with the model file and provenance when training a model, and the predictions and provenance when predicting GO terms.

The features and label files are used directly as an input to the AdaBoost model without scaling or normalizing, as AdaBoost does not benefit from this. AdaBoost is used with the following parameters: base_estimator=DecisionTreeClassifier, max_depth=5, algorithm=‘SAMME, learning_rate=1.0, n_estimators=100,000. AdaBoost scores are calibrated using SKLearn’s CalibratedClassifierCV using the following parameters: cv=prefit, method=sigmoid,

When using ‘CrowdGO_train.py’ the model file is written to the output folder as ‘model.pkl’, and when using CrowdGO.py’ the predictions are written to the output folder as ‘crowdgo.tab’ and ‘crowdgo_raw.csv’ as the direct raw output of ADABoost for each input row.

## Acknowledgements

The authors thank Romain Feron, Livio Ruzzante, and Antonin Thiébaut for providing useful suggestions for improvements and valuable feedback on the manuscript.

