## Supplementary data for "CrowdGO: machine learning and semantic similarity guided consensus Gene Ontology annotation"

### *Supplementary Information*

**Table S1. Checklist for reporting and evaluating machine learning models**

Based on *Best practices in machine learning for chemistry*, Artrith et al. Nature Chemistry vol. 13, pg 505–508, 2021, https://doi.org/10.1038/s41557-021-00716-z

| | |
|---|---|
| **1. Data sources** | |
| 1a. Are all data sources listed and publicly available? | All proteins used for the evaluation of CrowdGO in Figure 1, Figure 2, and Table 2 are available in UniProt and the GOA database.<br><br>All proteins used for the evaluation of Table 3 are taken from the CAFA3 challenge data set found at, amongst others, https://github.com/ashleyzhou972/CAFA_assessment_tool<br><br>All proteins used for the evaluation of Figure 3 are taken from UniProt.<br><br>Details are listed under the ‘Proteins and gene ontology annotations’ subsection of the ‘Materials and Methods’ section. |
| 1b. If using an external database, is an access date or version number provided? | CrowdGO uses several external databases. All versions used for the evaluation in Figure 1, Figure 2, Table 2, and Table 3 are found under the ‘Running input predictors and CrowdGO’ subsection of the ‘Materials and Methods’ section.<br><br>Additionally, Figure 3 uses the trained models on the data listed above, but uses different data for predicting GO terms. These can be found under the ‘Annotating model and non-model species’ subsection of the ‘Materials and Methods’ section. |
| 1c. Are any potential biases in the source dataset reported and/or mitigated? | All data used for the training and evaluation of CrowdGO, including its input predictors as part of its pipeline, are from before the data used for evaluation and thus no bias is introduced. The exception is the data used for Figure 3, but because this figure doesn’t evaluate against a test set and instead uses a subjective evaluation bias is no factor here. |
| **2. Data cleaning** | |
| 2a. Are the data cleaning steps clearly and fully described, either in text or as a code pipeline? | Most data cleaning steps as part of the CrowdGO pipeline used for the evaluation in this paper are described on the gitlab wiki https://gitlab.com/mreijnders/CrowdGO. |

| | |
|---|---|
| | Additionally the training and test set is cleaned as described in the 'Proteins and gene ontology annotations' subsection of the 'Materials and Methods' section. |
| 2b. Is an evaluation of the amount of removed source data presented? | A report of the numbers of proteins removed from the training and test set is described in the 'Proteins and gene ontology annotations' subsection of the 'Materials and Methods' section. |
| 2c. Are instances of combining data from multiple sources clearly identified, and potential issues mitigated? | The paper describes a methodology on combining predictions from different sources and is elaborately described throughout. |
| **3. Data representations** | |
| 3a. Are methods for representing data as features or descriptors clearly articulated, ideally with software implementations? | Features used and a description of how the features are gathered are described under the 'The CrowdGO Algorithm' subsection of the 'Materials and Methods' section. The gathering of these features is available as part of the CrowdGO pipeline and software package. |
| 3b. Are comparisons against standard feature sets provided? | Not applicable |
| **4. Model choice** | |
| 4a. Is a software implementation of the model provided such that it can be trained and tested with new data? | All models described in the paper are available at https://gitlab.com/mreijnders/CrowdGO |
| 4b. Are baseline comparisons to simple/trivial models (for example, 1-nearest neighbour, random forest, most frequent class) provided? | CrowdGO is a predictor that combines existing predictors to improve predictions. These input predictors are thus provided as baselines to demonstrate CrowdGO does indeed improve over these predictors. |
| 4c. Are baseline comparisons to current state-of-the-art provided? | The CAFA3 top performers are used to compare against CrowdGO on the CAFA3 challenge set.<br><br>Additionally, several of the methods used as an input to CrowdGO, which are used as baseline comparisons as described in section 4b, are considered state-of-the-art. |
| **5. Model training and evaluation** | |
| 5a. Does the model clearly split data into different sets for training (model selection), validation (hyperparameter optimization), and testing (final evaluation)? | No hyperparameter optimization is performed.<br><br>The training and test set are split 50/50, based on proteins, as described in the 'Proteins and gene ontology annotations' subsection of the 'Materials and Methods' section. Additionally, the CAFA3 challenge data is used for testing, therefore these proteins are not used as part of the training set.<br><br>Table 1 provides an overview of all data sets used for training and testing. |
| 5b. Is the method of data split (data splitting (for example, random, cluster- or time-based splitting, forward cross-validation) clearly | Data splitting for the training and test set is described in the 'Proteins and gene ontology annotations' subsection of the 'Materials and Methods' section.<br><br>Otherwise not applicable. |

| | |
|---|---|
| stated? Does it mimic anticipated real-world application? | |
| 5c. Does the data splitting procedure avoid data leakage (for example, is the same composition present in the training and test sets)? | There is no overlap between the training and test set, as is described in the ‘Proteins and gene ontology annotations’ subsection of the ‘Materials and Methods’ section. |
| **6. Code and reproducibility** | |
| 6a. Is the code or workflow available in a public repository? | Yes: https://gitlab.com/mreijnders/CrowdGO |
| 6b. Are scripts to reproduce the findings in the paper provided? | Yes: https://gitlab.com/mreijnders/CrowdGO under the ‘supplementary’ section. |

### Supplementary Figures S1, S2, and S3.

**Figure S1: Distributions of GO terms per protein for the V162-V198 training and testing datasets and the CAFA3 benchmarking dataset.**

Distributions are shown as density plots (A-C) and as empirical cumulative distribution function (ECDF) plots (D-E) for the Molecular Function (MFO), Biological Process (BPO), and Cellular Component (CCO) ontologies. The total number of proteins in each dataset are shown on the plots, along with results from Wilcoxon (Mann-Whitney) tests for each pair of datasets for A-C, and with results of Kolmogorov-Smirnov tests for each pair of datasets for D-F. Analysis, plotting, and significance testing all performed using R 3.6.1. The V162-V198 training and testing datasets show no significant differences in their distributions of GO terms per protein. Compared with the CAFA3 dataset, they are both significantly lower for MFO, significantly higher for BPO, and not significantly different for CCO.

**Figure S2: Distributions of information contents for GO terms from the V162-V198 training and testing datasets and the CAFA3 benchmarking dataset.**
Distributions are shown as density plots (A-C) and as empirical cumulative distribution function (ECDF) plots (D-E) for the Molecular Function (MFO), Biological Process (BPO), and Cellular Component (CCO) ontologies. The total number of gene-term annotations in each dataset are shown on the plots, along with results from Wilcoxon (Mann-Whitney) tests for each pair of datasets for A-C, and with results of Kolmogorov-Smirnov tests for each pair of datasets for D-F. Analysis, plotting, and significance testing all performed using R 3.6.1. The V162-V198 training and testing datasets show no significant differences in their distributions of information contents (ICs). Compared with the CAFA3 dataset, their medians are not significantly different for MFO but their distributions are significantly flatter (the peak in the CAFA3 dataset arises from GO:0042802 ‘identical protein binding’ with an IC of 6.5 annotated to 61 proteins, the most proteins for any term). Their ICs are significantly higher than the CAFA3 dataset for both BPO and CCO.

**Figure S3: CrowdGO applied to functionally annotate the complete proteomes of taxonomically diverse species.**
Results for eight species are shown in main text Figure 3, here results for the remaining four species are presented. CrowdGO consensus annotation results were compared with existing Gene Ontology (GO) term annotations from UniProt, and with the subsets of manually curated SwissProt (where available) and automatically inferred TrEMBL annotations, for (A) *Pan troglodytes* Chimpanzee and *Homo sapiens* human, (B) *Arabidopsis thaliana* the thale cress (repeated here for plant-plant model-non-model comparison) and *Solanum lycopersicum* Tomato, and (C) *Candidatus Thorarchaeota archaeon* SMTZ1-45. Bars at the top of each panel show the total numbers of proteins annotated with at least one GO term for each respective annotation dataset, with white-filled areas showing the remaining proteins with no annotations. Split violin plots show the distributions of the numbers of GO Slim terms annotated per protein (purple, left), and summed leaf+parents information content (IC) per protein (green, right). Y-axes are limited to a maximum of value of 300 for total protein IC distributions. The boxplots show the medians and 1.5 times the interquartile range for the numbers of GO Slim terms annotated per protein.

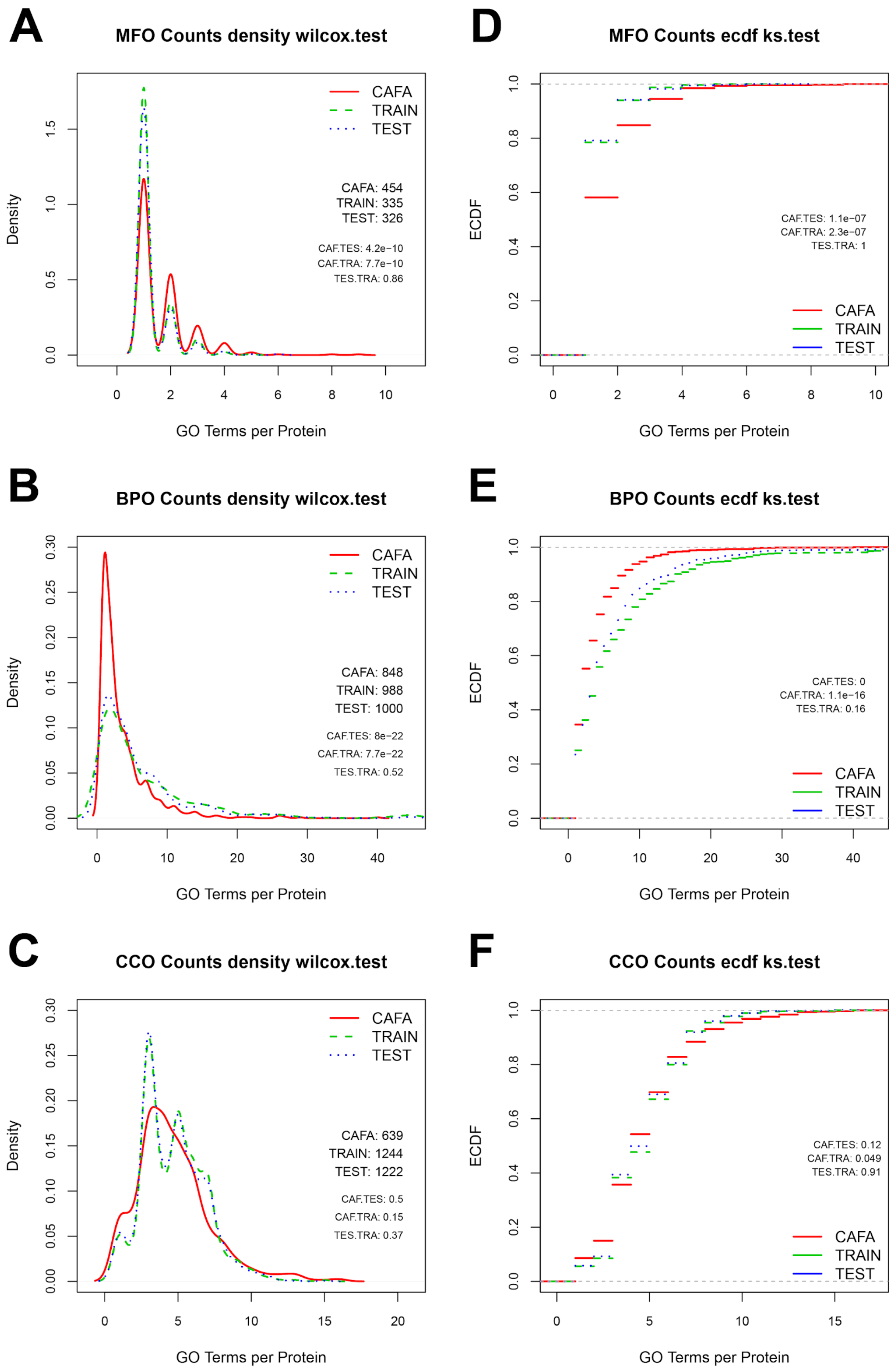
A
MFO Counts density wilcox.test
CAFA
TRAIN
TEST
CAFA: 454
TRAIN: 335
TEST: 326
CAF.TES: 4.2e−10
CAF.TRA: 7.7e−10
TES.TRA: 0.86
Density
GO Terms per Protein
D
MFO Counts ecdf ks.test
CAF.TES: 1.1e−07
CAF.TRA: 2.3e−07
TES.TRA: 1
ECDF
GO Terms per Protein
B
BPO Counts density wilcox.test
CAFA: 848
TRAIN: 988
TEST: 1000
CAF.TES: 8e−22
CAF.TRA: 7.7e−22
TES.TRA: 0.52
E
BPO Counts ecdf ks.test
CAF.TES: 0
CAF.TRA: 1.1e−16
TES.TRA: 0.16
C
CCO Counts density wilcox.test
CAFA: 639
TRAIN: 1244
TEST: 1222
CAF.TES: 0.5
CAF.TRA: 0.15
TES.TRA: 0.37
F
CCO Counts ecdf ks.test
CAF.TES: 0.12
CAF.TRA: 0.049
TES.TRA: 0.91

**Figure S1**

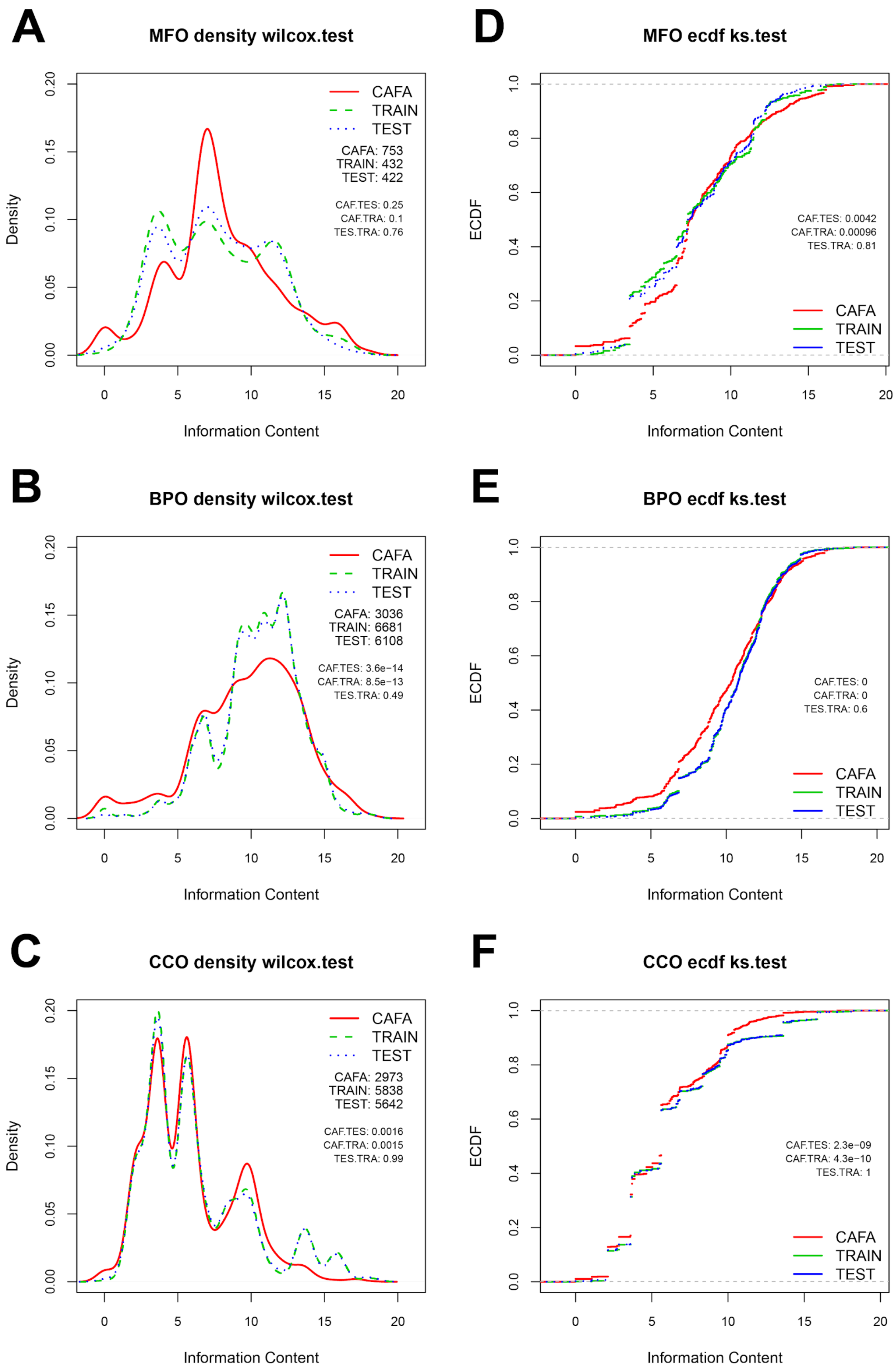
A
MFO density wilcox.test
Density
CAFA
TRAIN
TEST
CAFA: 753
TRAIN: 432
TEST: 422
CAF.TES: 0.25
CAF.TRA: 0.1
TES.TRA: 0.76
Information Content
D
MFO ecdf ks.test
ECDF
CAF.TES: 0.0042
CAF.TRA: 0.00096
TES.TRA: 0.81
CAFA
TRAIN
TEST
Information Content
B
BPO density wilcox.test
Density
CAFA
TRAIN
TEST
CAFA: 3036
TRAIN: 6681
TEST: 6108
CAF.TES: 3.6e−14
CAF.TRA: 8.5e−13
TES.TRA: 0.49
Information Content
E
BPO ecdf ks.test
ECDF
CAF.TES: 0
CAF.TRA: 0
TES.TRA: 0.6
CAFA
TRAIN
TEST
Information Content
C
CCO density wilcox.test
Density
CAFA
TRAIN
TEST
CAFA: 2973
TRAIN: 5838
TEST: 5642
CAF.TES: 0.0016
CAF.TRA: 0.0015
TES.TRA: 0.99
Information Content
F
CCO ecdf ks.test
ECDF
CAF.TES: 2.3e−09
CAF.TRA: 4.3e−10
TES.TRA: 1
CAFA
TRAIN
TEST
Information Content

**Figure S2**

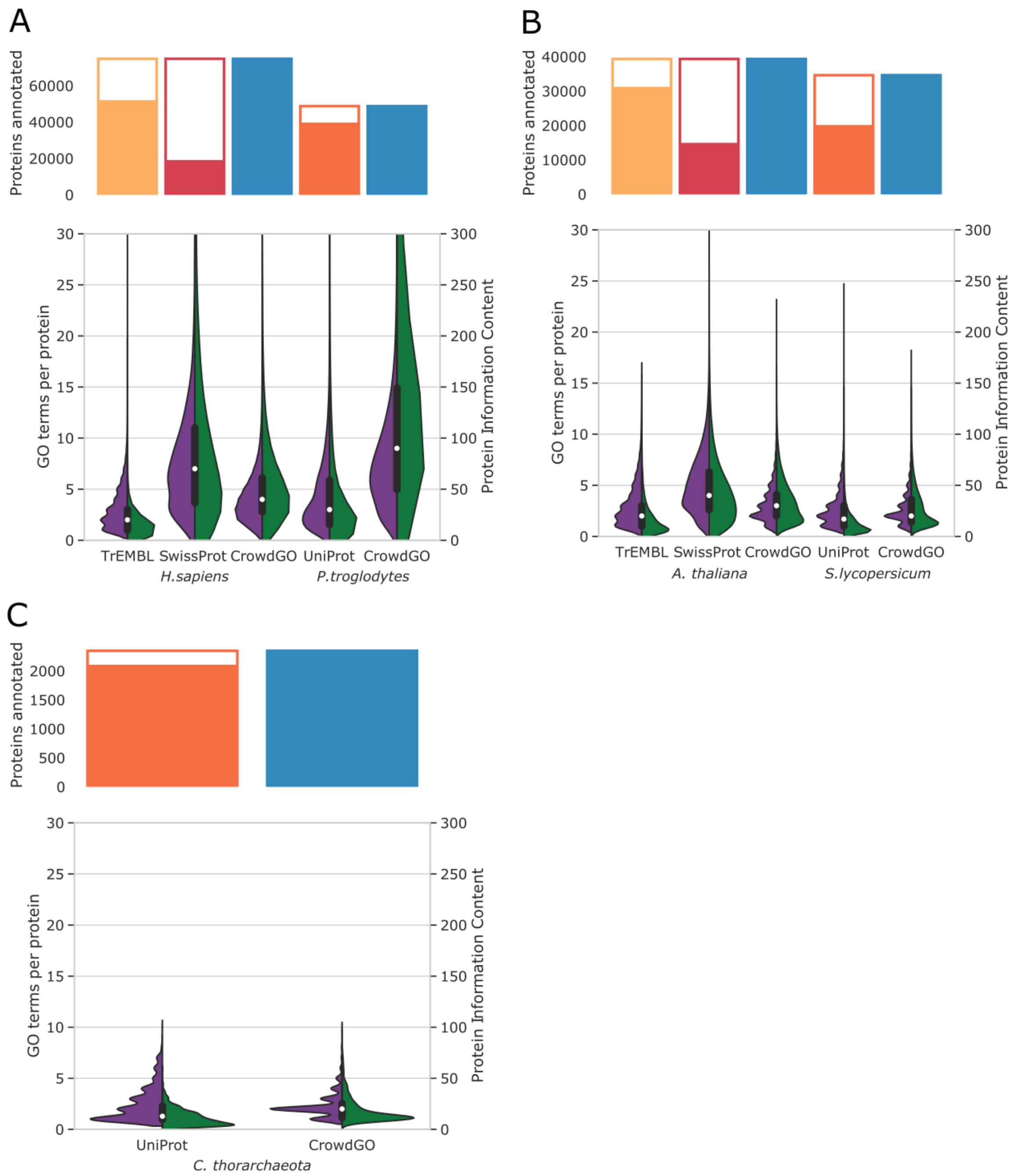
A
Proteins annotated
0
20000
40000
60000
GO terms per protein
Protein Information Content
TrEMBL
SwissProt
CrowdGO
UniProt
CrowdGO
H.sapiens
P.troglodytes
B
A. thaliana
S.lycopersicum
C
C. thorarchaeota

**Figure S3**
